# Reconstructing the genetic relationship between ancient and present-day Siberian populations

**DOI:** 10.1101/2023.08.21.554074

**Authors:** Haechan Gill, Juhyeon Lee, Choongwon Jeong

**Affiliations:** School of Biological Sciences, Seoul National University, Seoul, 08826, Republic of Korea

## Abstract

Human populations across a vast area in northern Eurasia, from Fennoscandia to Chukotka, share a distinct genetic component often referred to as the Siberian ancestry. Most enriched in present-day Samoyedic-speaking populations such as Nganasans, its origins and history still remain elusive despite the growing list of ancient and present-day genomes from Siberia. Here we reanalyze published ancient and present-day Siberian genomes focusing on the Baikal and Yakutia, resolving key questions regarding their genetic history. First, we show a long-term presence of a unique genetic profile in southern Siberia, up to 6,000 years ago, which distinctly shares a deep ancestral connection with Native Americans. Second, in the Baikal we find no direct contribution of the Early Neolithic Kitoi people to Late Neolithic and Early Bronze Age Serovo-Glazkovo ones. Third, the Middle Neolithic individual from Yakutia, belonging to the Belkachi culture, serves as the best source so far available for the spread of the Siberian ancestry into Fennoscandia and Greenland. These findings shed light on the genetic legacy of the Siberian ancestry and provide insights into the complex interplay between different populations in northern Eurasia throughout history.

## Introduction

Migration and admixture are key demographic events that have influenced the genetic structure of modern human populations (Patterson et al., 2012). The genetic diversity of inhabitants in Inner Eurasia, a vast geographic region encompassing Siberia and the Eurasian Steppe, has been shaped by a complex history of mixture between diverse source populations of both eastern and western Eurasian origins (Jeong et al., 2019). As a result of this complex history, present-day Inner Eurasian populations are stratified into three distinct admixture clines mirroring geography. The northernmost one among these clines, composed of populations from the boreal forest and tundra regions who mostly speak the Uralic and Yeniseian languages, share a distinct type of Eastern Eurasian ancestry, frequently referred to as the Siberian ancestry in recent archaeogenetics literature (Tambets et al., 2018). Among the present-day populations, it is most enriched in Nganasans and other Samoyedic-speaking ones such as Nenets, Enets, and Selkups.

The Samoyedic-speaking populations inhabit the northernmost region of Siberia (Nganasans, Enets, Nenets) as well as the Yenisei River basin to the south (Selkups). Although ancient genomes have been only scarcely reported in these regions, the Siberian ancestry was present in a larger area in the past, including early Metal Age individuals from Bolshoy Oleni Ostrov in the Kola Peninsula, Iron Age individuals from the eastern Baltic Sea, and Iron Age individuals from the Volga-Oka interfluve (Lamnidis et al., 2018; Saag et al., 2019; Peltola et al., 2023). Together with the genetic analysis of present-day populations, these studies suggest that the populations with the Siberian ancestry once occupied a large area in Siberia and northeastern Europe and formed a substratum for the genetic profile of present-day populations in the regions. Therefore, tracing the origins and spreads of the Siberian ancestry is crucial for understanding the formation of present-day human populations and languages in northern Eurasia. Despite the recent accumulation of ancient genome data in Siberia, centered on southern Siberia, it remains obscure how these populations are related to present-day inhabitants of Siberia, such as Nganasan, calling for a careful re-investigation of previously published data.

Among the previously published ancient genomes, the Middle Holocene (ca. 8,000-3,000 years ago) southern Siberians represent a pivotal ancient lineage connected to the present-day Siberian ancestry (Sandweiss et al., 1999). Of particular interest are those from the archaeological sites at Lake Baikal and Yakutia because they harbored Y haplogroups N and Q, which are prevalent in present-day Siberians (Karafet et al., 2018). Overall, they show genome-wide genetic profiles closely related to present-day Siberians but still different enough to reject a simple ancestor-descendant relationship. Therefore, the Middle Holocene Siberians in Yakutia and Lake Baikal provide an excellent starting point from which we can build a historical model for the origins of the Siberian ancestry and populations harboring it.

The genetic makeup of the Middle Holocene Siberians resulted from the admixture of three ancestries (Figure 1): Ancient North Eurasian (ANE), Ancient Paleo-Siberian (APS; an ancestry closely related to the Native American ancestries), and Ancient North Asian (ANA). ANE ancestry is represented by the Upper Paleolithic individuals from the Mal’ta (MA1) and Afontova Gora sites (AG2 and AG3) (Raghavan et al., 2014; Fu et al., 2016). During the Last Glacial Maximum (LGM), ANE ancestry intermixed with populations of Eastern Eurasian origin and formed the ancestral population of Native Americans. This ancestral population left its genetic legacy in later APS populations, e.g. 14,000-year-old terminal Pleistocene individual from Ust-Kyakta-3 in southern Siberia (UKY) and 9,800-year-old Mesolithic individual from the Duvanny Yar site at the Kolyma River in northern Siberia (Kolyma_M) (Sikora et al., 2019; Yu et al., 2020). At the beginning of the Middle Holocene, individuals of ANA ancestry already appeared in both East and West Baikal (Kilinc et al., 2021), presumably expanded from the neighboring regions in northeastern China where ANA ancestry were present at least 14,000 years ago (Siska et al., 2017; Ning et al., 2020; Mao et al., 2021). Although the genetic profile and the geographic distribution of these three ancestries in Siberia have been actively investigated using ancient genomes, it remains unexplored how, when, and where the genetic profiles of present-day Siberians were formed out of these three ancestral populations. In this study, we provide a proximal historical model for the genetic relationship between a comprehensive set of published Siberian genome data. Our findings demonstrate that the APS population was present in the Baikal and Yakutia during the Middle Holocene and in each region the local APS population formed a genetic substratum for the later populations. Furthermore, in the Baikal we find no direct genetic contribution from the Early Neolithic Kitoi populations to the Late Neolithic and Early Bronze Age Serovo-Glazkovo ones, resolving a long-standing archaeological question about their relationship. Finally, we show that the Middle Neolithic Yakutia population serves as a better-fitting source than the Lake Baikal one for the Siberian ancestry found in the genetic makeup of northern Siberians, northeastern Europeans, Paleo-Eskimos, and ancient Athabaskans.

**Figure 1.**
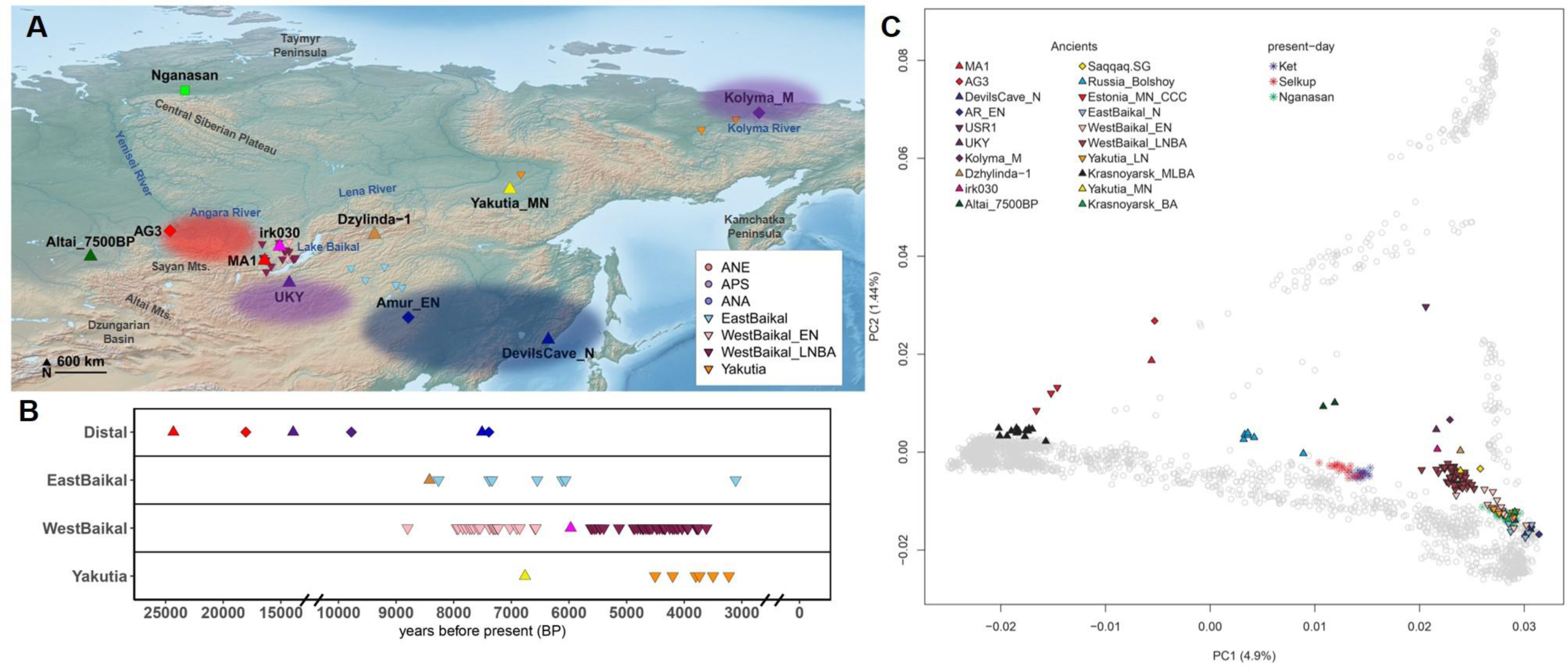
**(A) Geographic sites of analyzed samples in this study.** Three distal source populations (AG3, MA1 for ANE; UKY, Kolyma_M for APS; Amur_EN, DevilsCave_N for ANA), which are used for explaining the formation of Middle Holocene Siberians, are represented in red, purple, and blue. Their virtual occupancies are presented by transparent ellipses. Among Middle Holocene Siberians, key individuals, irk030, Dzhylinda-1, Altai_7500BP, and Yakutia_MN, are marked separately, and the rest of the Middle Holocene and present-day populations are marked based on geographic locations and archaeological period. The base map data is downloaded from Natural Earth (https://www.naturalearthdata.com/downloads/), and the coordinates of each sample are from the AADR annotation file v52.2. **(B) The radiocarbon dating of three distal sources and Middle Holocene Siberians.** The dating results are from AADR annotation file v52.2 and presented in before the present (BP). **(C) Principal component analysis performed with present-day Eurasian and American individuals.** Each present-day sample is placed on principal component 1 and 2 coordinates by grey circles. Key ancient and present-day genomes are projected on pre-calculated principal components and labeled.

## Results

### The genetic profile of ancient Siberian individuals

We curated published genomes and genome-wide data of ancient Siberian populations focusing on Lake Baikal and Yakutia. Most ancient individuals date to the Middle Holocene, ranging 8,800-3,100 BP (Figure 1). To overview the genetic profile of these ancient individuals, we performed principal component analysis (PCA) (Patterson et al., 2006) and projected them onto the principal components (PCs) calculated from 2,270 present-day Eurasian and American individuals (Figure 1). PC1 separates individuals from west to east, and PC2 separates from Eurasians to Native Americans. While most Middle Holocene Siberian individuals fall on a cline in the PC space between the ANE and ANA populations, two individuals deviate from this cline: Dzhylinda-1, the earliest individual among east Baikal individuals (6,564-6,429 cal. BCE), and irk030, the earliest Late Neolithic west Baikal individual (4,150-3,950 cal. BCE). They are shifted upward along PC2, suggesting an extra affinity with Native Americans.

For the group-based analyses, we removed PCA outliers and 1st-degree relatives and allocated the ancient Siberian individuals into six analysis super-groups according to their geographic location, archaeological period, and PCA pattern: EastBaikal_M, EastBaikal_N, WestBaikal_EN, WestBaikal_LNBA, Yakutia_MN, and Yakutia_LN (de Barros Damgaard et al., 2018; Sikora et al., 2019; Yu et al., 2020; Kılınç et al., 2021) (Table S1).

### The genetic legacy of the ancestral Native American gene pool in Holocene Siberia

We first model the genetic profile of the two ancient Siberian individuals, Dzhylinda-1 and irk030, who potentially show a link to the ancestral Native American gene pool (Figure 1). Of note, two earlier-period Ancient Paleo-Siberian individuals, 14,000-year-old UKY and 9,800-year-old Kolyma_M, show a similar shift in the PC space toward Native Americans to a greater degree. We formally tested if an ancestry component related to Native Americans is required to explain the genetic profile of these Siberian individuals using *qpAdm* (Lazaridis et al., 2016). While the two-way admixture model of ANE+ANA does not fit them, the three-way model of ANE+ANA+Native American adequately fits them with similar ancestry proportions with the APS individuals (Figure 2, Table S2). Indeed, using *qpWave* (Reich et al., 2012) we show that (UKY, Kolyma_M, irk030) and (UKY, Kolyma_M, Dzhylinda-1) can be modeled as a clade, respectively (Table S2). However, Dzhylinda-1 and irk030 do not form a clade in *qpWave* analysis, suggesting a difference in the admixture proportions between these two Holocene individuals.

**Figure 2.**
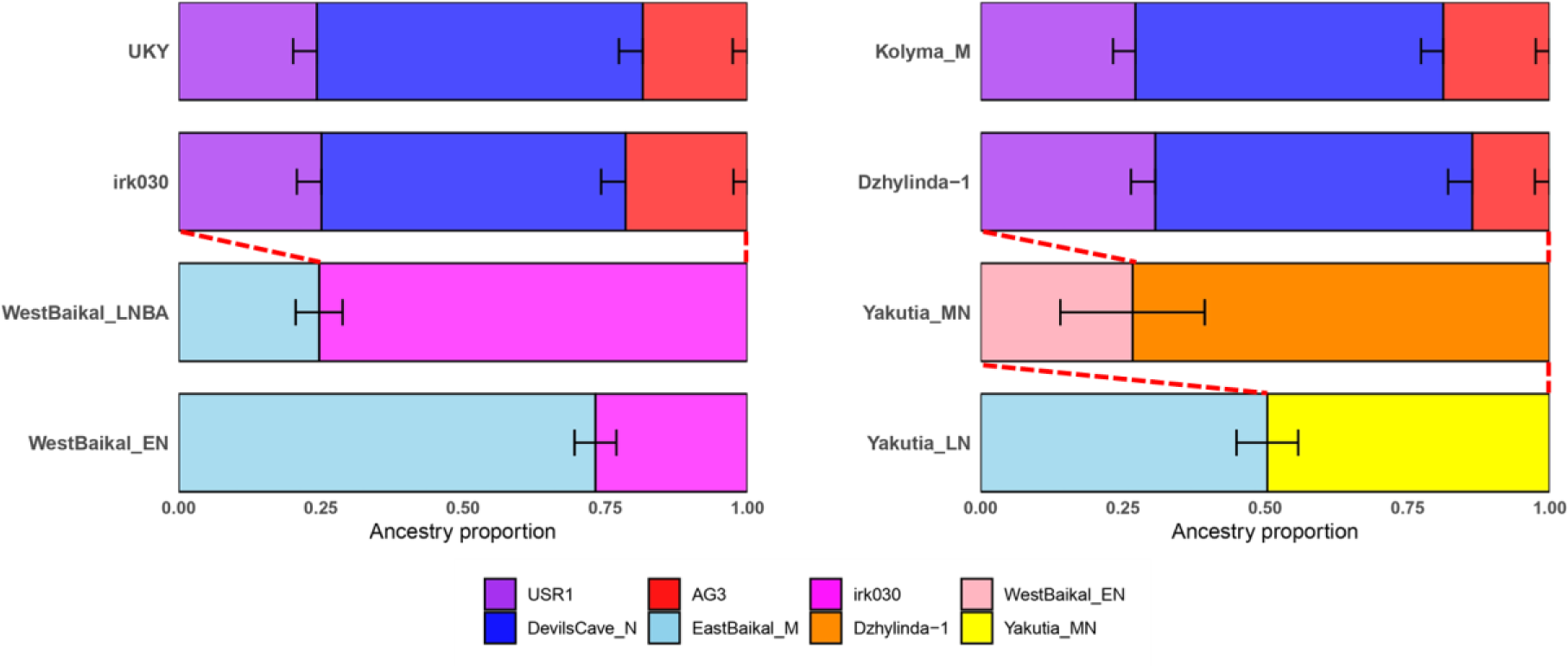
The genetic profiles of ancient Siberians estimated by *qpAdm*. Target populations are modeled as a mixture of two or three populations. The ancestry proportion of each source is represented by the box size on the x-axis. Horizontal bars represent ±1 SE estimated by *qpAdm* using 5cM block jackknifing. Detailed results are presented in Table S2 and S3.

A recent study reported a distinctive gene pool of Altai hunter-gatherers (Altai_HG) who were modeled to carry a combination of the ANE and APS ancestries (Wang et al., 2023). We found that Altai_HG is adequately modeled as irk030/Dzhylinda-1 + ANE, suggesting that they are good proxies of the APS ancestry in Altai_HG (Table S2). Interestingly, irk030 is found to be the best APS proxy for Altai_HG by *qpAdm* rotating approach (Harney et al., 2021) (Table S2). Together with the geographic and chronological proximity, we propose that APS ancestry continued from the Upper Paleolithic (UKY) to at least the Late Neolithic (irk030) in Siberia and mixed with a local ANE population in the Altai mountains further to the west.

By utilizing outgroup-*fз* statistics in the form of *fз*(Mbuti; irk030/Dzhylinda-1, X), we searched for the genetic link of irk030 and Dzhylinda-1 with later populations (Figure S2, Table S6). Dzhylinda-1 and irk030 showed a high genetic affinity with Middle Neolithic Yakutia (Yakutia_MN) and Late Neolithic-Bronze Age West Baikal population (WestBaikal_LNBA), respectively. *F₄* statistics in the form of *f₄* (Mbuti, irk030 /Dzhylinda-1; WestBaikal_LNBA, Yakutia_MN) confirm that Dzhylinda-1 and irk030 are closer to Yakutia_MN and WestBaikal_LNBA, respectively (Figure S3, Table S5). These findings suggest that the difference between the Yakutia and west Baikal populations trace back to the distinct Middle Holocene APS populations represented by Dzhylinda-1 and irk030, respectively. We note that Dzhylinda-1 is from the northern part of the East Baikal region, close to but not in Yakutia, but utilize him for modeling Yakutian populations based on this genetic affinity.

### Discontinuity between the early Neolithic and later populations in West Baikal

The genetic profile of the West Baikal populations underwent a transition from ANA-rich Early Neolithic West Baikal population (WestBaikal_EN) to ANE-rich WestBaikal_LNBA, coinciding with the cultural shift from the Kitoi to the Serovo-Glazkovo culture (Weber et al., 2002). Previous studies have suggested that the resurgence of the ANE ancestry in Serovo-Glazkovo was due to gene flow from an APS source like UKY, Kolyma_M, and Altai_HG, assuming the local Kitoi culture as a substratum (Sikora et al., 2019; Yu et al., 2020; Wang et al., 2023). However, differences in major Y haplogroup, subsistence strategy, social structure, and cranial morphology between the two cultures, along with a significant timing gap of approximately 800 years, raise the possibility of discontinuity (Weber et al., 2002; Movsesian et al., 2014; Kılınç et al., 2021). This implies an alternative scenario that the Serovo-Glazkovo population formed elsewhere without a direct contribution from the Kitoi one and replaced it. We focused on irk030, the oldest individual from the Serovo-Glazkovo culture, and examined its genetic affinity with APS as well as its relationship with other West Baikal populations. Using *f₄* symmetry test, we found that irk030 has higher affinity with the ANE ancestry represented by AG3 and Tarim_EMBA1, while other West Baikal populations, both the Early Neolithic Kitoi and the later Serovo-Glazkovo ones, have more ANA ancestry represented by East Baikal populations (Figure S3, Table 1, Table S5). We further explored the genetic makeup of the West Baikal populations by investigating the admixture model between ANA and irk030 and found EastBaikal_M+irk030 adequately fit both West Baikal populations (Figure 2, Table S3). We replicate previous reports on the difference between the genetic makeup of WestBaikal_EN and WestBaikal_LNBA: WestBaikal_EN showed a relatively lower contribution from irk030 (p=0.610; 27% from irk030), while WestBaikal_LNBA showed a higher contribution from irk030 (p=0.333; 75% from irk030) (Figure 2, Table 1, Table S3). Interestingly, using the *qpAdm* rotating approach we show that the ANA ancestry in WestBaikal_LNBA is better modeled by EastBaikal_N than by WestBaikal_EN (Table S3). Taken together, our findings support an alternative explanation for the genetic makeup of the Serovo-Glazkovo culture, emphasizing the significant contribution of APS population and ANA-related gene flow originated from East Baikal rather than the Kitoi culture.

**Table 1.** Key genetic symmetricity test and admixture modeling results. (A) *f*4(pop1, pop2; pop3, pop4) for key genetic tests. Z-scores were calculated by dividing f4 by the standard error measure (s.e.m.) estimated by 5 cM block-jackknifing. (B) Key qpAdm and qpWave results. Coeff1 and coeff2 represent the ancestry proportion (± 1 s.e.m.) contributed by ref1 and ref2 populations, respectively. “Table” column indicates the supplementary table that includes the result.

| **A. Key $f_4$ statistics results between target populations.** |  |  |  |  |  |  |
| --- | --- | --- | --- | --- | --- | --- |
| **pop1** | **pop2** | **pop3** | **pop4** | **$f_4$** | **Z-score** | **Table** |
| Mbuti | irk030 | WestBaikal_LNBA | Yakutia_MN | -0.00187 | -3.833 | S5B |
| Mbuti | Dzhylidna-1 | WestBaikal_LNBA | Yakutia_MN | 0.00190 | 3.256 | S5B |
| Mbuti | EastBaikal_N | irk030 | WestBaikal_EN | 0.00487 | 13.032 | S5D |
| Mbuti | EastBaikal_N | irk030 | WestBaikal_LNBA | 0.00239 | 6.557 | S5C |
| Mbuti | WestBaikal_EN | Dzhylinda-1 | Yakutia_MN | 0.00161 | 3.356 | S5E |
| Mbuti | EastBaikal_N | Yakutia_MN | Yakutia_LN | 0.00238 | 5.525 | S5F |
| Mbuti | Krasnoyarsk_MLBA | Yakutia_MN | Selkup | 0.00199 | 5.464 | S5H |
| Mbuti | Krasnoyarsk_MLBA | Yakutia_MN | Ket | 0.00162 | 4.599 | S5J |
| Mbuti | Estonia_MN_CCC | Yakutia_MN | Russia_Bolshoy | 0.00770 | 10.444 | S5K |
| **B. Key qpAdm and qpWave results of target populations.** |  |  |  |  |  |  |
| **target** | **ref1** | **ref2** | **p-value** | **coeff1** | **coeff2** | **Table** |
| irk030 | UKY | - | 1.00E-01 | - | - | S2 |
| WestBaikal_EN | irk030 | EastBaikal_M | 6.15E-01 | 0.268±<br>0.037 | 0.732±<br>0.037 | S3 |
| WestBaikal_LNBA | irk030 | EastBaikal_N | 1.36E-01 | 0.759±<br>0.040 | 0.241±<br>0.040 | S3 |
| Dzhylinda-1 | UKY | - | 7.88E-01 | - | - | S2 |
| Yakutia_MN | Dzhylinda-1 | WestBaikal_EN | 2.07E-01 | 0.732±<br>0.125 | 0.268±<br>0.125 | S3 |
| Yakutia_LN | Yakutia_MN | EastBaikal_N | 2.90E-01 | 0.355±<br>0.042 | 0.645±<br>0.042 | S3 |
| Krasnoyarsk_BA | Yakutia_MN | EastBaikal_N | 8.48E-01 | 0.475±<br>0.094 | 0.525±<br>0.094 | S4 |
| Nganasan | Yakutia_MN | EastBaikal_N | 7.96E-01 | 0.595±<br>0.063 | 0.405±<br>0.063 | S4 |
|  | Krasnoyarsk_BA | - | 1.94E-01 | - | - | S4 |
| Selkup | Yakutia_MN | Krasnoyarsk_MLBA | 7.18E-02 | 0.726±<br>0.013 | 0.274±<br>0.013 | S4 |
| Ket | Yakutia_MN | Krasnoyarsk_MLBA | 1.38E-01 | 0.771±<br>0.014 | 0.229±<br>0.014 | S4 |
| Russia_Bolshoy | Yakutia_MN | Estonia_MN_CCC | 4.53E-01 | 0.516±<br>0.019 | 0.484±<br>0.019 | S4 |
| Saqqaq | Yakutia_MN | - | 2.11E-01 | - | - | S4 |
| Athabaskan_1100BP | Yakutia_MN | Clovis | 2.64E-01 | 0.374±<br>0.029 | 0.626±<br>0.029 | S4 |

### Repeated introduction of ANA ancestry in Yakutia populations

In Yakutia, we first tested if Dzhylinda-1, Yakutia_MN, and Yakutia_LN form a simple time series of local population continuity without gene flow from other sources. Using *f₄* symmetry test, we found that WestBaikal_EN is closer to Yakutia_MN than its preceding Dzhylinda-1, and ANA-related populations are closer to Yakutia_LN than its preceding Yakutia_MN (Figure S3, Table S5). Formally modeling this relationship with *qpAdm*, we show that Yakutia_MN and Yakutia_LN are adequately modeled as Dzhylinda-1+WestBaikal_EN (p=0.330; 29% contribution from WestBaikal_EN) and Yakutia_MN+EastBaikal_N (p=0.890; 55% contribution from EastBaikal_N), respectively (Figure 2, Table 1, Tables S3).

Lastly, we comprehensively tested all proximal models of the Middle Holocene Siberian populations by graph-based analysis, *qpGraph* (Figure 3). We construct a basal graph including Mbuti, MA1, Western European hunter-gatherers (WHG), EastBaikal_N, and USR1, referring to the previous study (Yu et al., 2020). We added irk030 and Dzhylinda-1 as an independent mixture of Native American and ANA branches. Then, we modeled West Baikal and Yakutia populations as successors of irk030 and Dzhylinda-1, respectively. The proximal admixture models are well explained in the statistically feasible final admixture graph (worst Z = -2.77), though we caution that we did not perform a comprehensive search over possible graph topologies.

**Figure 3.**
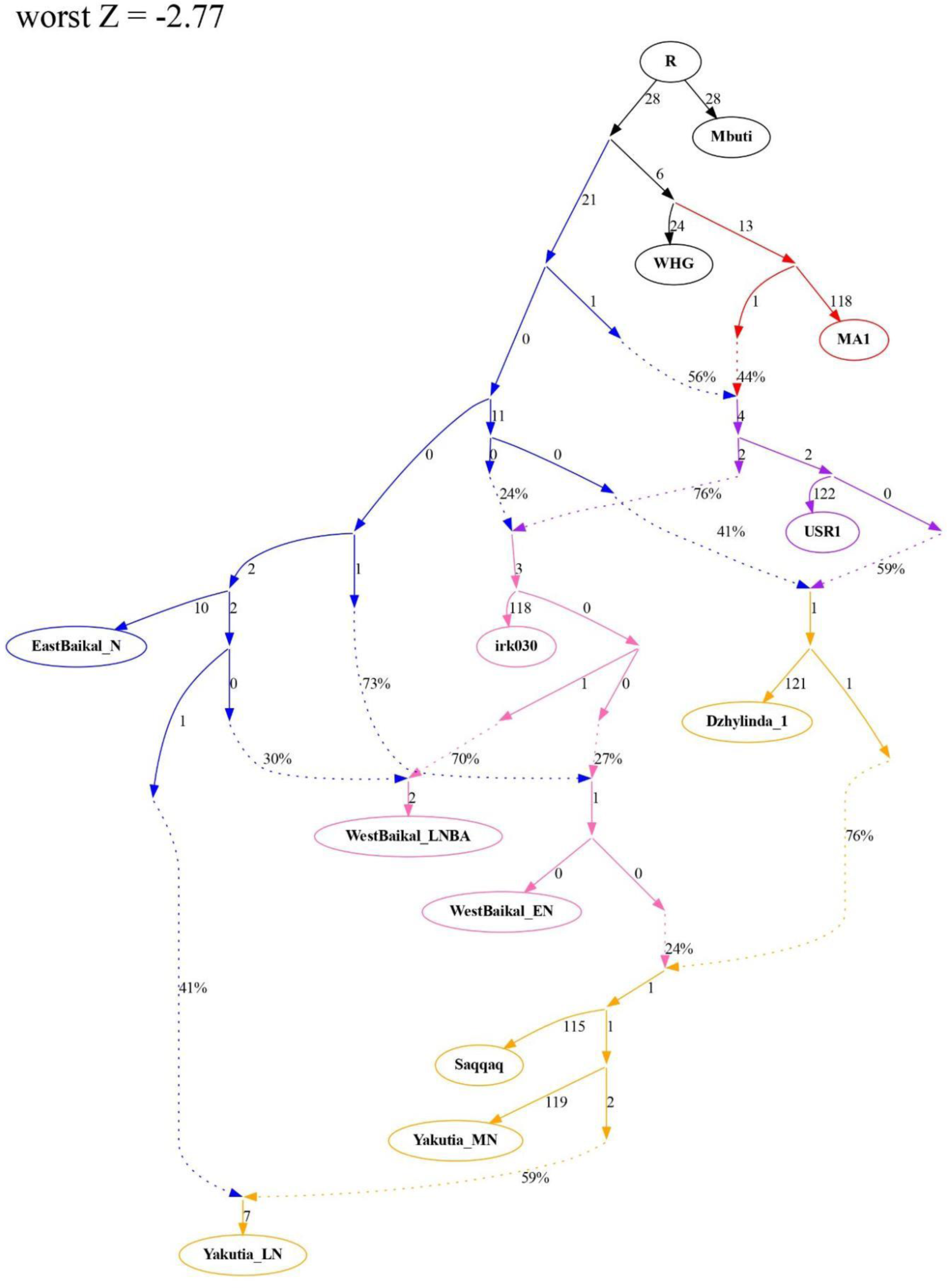
Manually fitted admixture graph explaining population dynamics of ancient Siberian populations. The admixture graph is based on proximal admixture modeling results and manually fitted by *qpgraph* function. ANE, ANA, Native American, WestBaikal, and Yakutia lineages are shown in red, blue, purple, light pink, and yellow, respectively. The worst Z score is -2.77.

### Siberian ancestry originated from Neolithic Yakutia populations

Our detailed examination of the Middle Holocene Siberian populations has provided significant insights into the origin of the Siberian ancestry. We identified two primary lineages within the Middle Holocene Siberians: the Lake Baikal lineage and the Yakutia lineage. Interestingly, we observed an increasing affinity between ancient Yakutia populations and present-day Nganasan over time, while this trend was not observed in the West Baikal populations (Figure S1). Further analysis using the outgroup-*f*₃ statistic revealed a strong genetic affinity between Nganasan and Yakutia, as well as a closely related individual in southern Siberia, Krasnoyarsk_BA (Figure S4, Table S6). Similar to Yakutia_LN and Krasnoyarsk_BA, Nganasan could be modeled by Yakutia_MN+EastBaikal_N (p=0.796; 60% contribution from Yakutia_MN), but its Yakutia_MN ancestry proportion was significantly higher than that of Yakutia_LN (45%; Figure 4, Table 1, Table S4). This model breaks when Yakutia_LN was added to the outgroup population set, supporting a strong affinity between Nganasan and Yakutia_LN (p=3.03×10^-11^; Table S4). Therefore, we suggest that Nganasan descended from a metapopulation to which Yakutia_LN and Krasnoyarsk_BA belonged but its direct ancestor had less contribution from the EastBaikal_N-related gene flow than Yakutia_LN.

**Figure 4.**
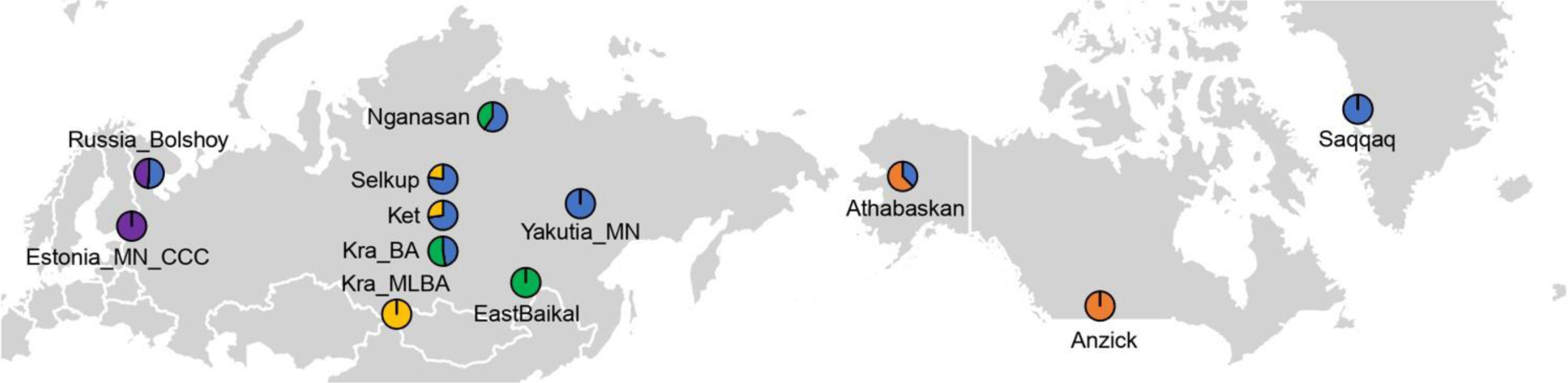
The genetic profiles of Siberian ancestry-related populations on a worldwide map. Yakutia_MN is used to represent Siberian ancestry, and each target is modeled as a mixture of Yakutia_MN and its local proximal source. All admixture models are feasible, and ancestry proportion is shown in pie charts (Table S4).

Other present-day Siberian populations, Selkups and Kets, also show high outgroup-*f*₃ statistics with ancient Yakutia populations (Figure S4). *F*₄ statistics in the form of *f₄*(Mbuti, X; Nganasan, Selkup) confirmed that ANE ancestry represented by AG3 and Eastern European hunter-gatherers (EHG) is necessary for Selkups (Figure S5, Table S5). However, both Nganasan+ANE and Yakutia_LN+ANE were infeasible for Selkups (Table S4). Hence, we used Yakutia_MN as a representative of Yakutia_LN+ANE, and then *f₄*(Mbuti, X; Yakutia_MN, Selkup) found an extra gene flow from the Bronze Age Steppe populations (Figure S5, Table 1, Table S5). Two-way admixture models, Yakutia_MN+Krasnoyarsk_MLBA, adequately explain both Selkups and Kets (p=0.072-0.140; 73% and 77% contribution from Yakutia_MN, respectively) (Figure 4, Figures S4-S5, Table 1, Table S4).

We tested whether the ancient Yakutia populations can represent Siberian ancestry first appeared in northeastern Europe, i.e. early Metal Age individuals in Bolshoy Oleni Ostrov (Russia_Bolshoy). Russia_Bolshoy has high outgroup*-f*₃ values with Tarim_EMBA1, Yakutia populations, and EHG (Figure S4, Table S6). We found that a parsimonious two-way admixture model, Yakutia_MN+Middle Neolithic Combed Ceramic Culture (Estonia_MN_CCC), is feasible (p=0.453; 52% contribution from Yakutia_MN), as well as a more complex three-way admixture model of Yakutia_LN+Estonia_MN_CCC+Tarim_EMBA1 (p=0.217; 39% contribution from Yakutia_LN, 16% contribution from Tarim_EMBA1, and 45% contribution from Estonia_MN_CCC) (Figure 4, Table 1, Table S4). Although less parsimonious, we still present the latter model for Y haplogroup information: Yakutia_LN and Russia_Bolshoy share Y haplogroup N, while Yakutia_MN belongs to Y haplogroup Q. Since only one individual is available, we cannot ignore the possibility that Y haplogroup N existed in the Middle Neolithic Yakutia population. Thus, further sampling on Middle Neolithic Yakutia is required to clarify the origin of Y haplogroup N in Russia_Bolshoy.

Regarding the origins of Paleo-Eskimo in Greenland, a cultural similarity with Belkachi, the Middle Neolithic Yakutia hunter-gatherer culture, has been suggested (Powers & Jordan, 1990; Coutouly, 2016; Flegontov et al., 2019; Kılınç et al., 2021). However, the Belkachi connection of Paleo-Eskimo has hardly been studied with ancient genome data. We confirm that Yakutia_MN, who belonged to the Belkachi culture, is cladal with a Paleo-Eskimo individual Saqqaq (qpWave p=0.211; Table 1, Table S4). In addition, Yakutia_MN and Saqqaq are exchangeable in ancestry modeling: Yakutia_MN+Anzick fits ancient Athabaskans (p=0.264; 37% contribution from Yakutia_MN) and Saqqaq+Krasnoyarsk_MLBA adequately fits Ket (p=0.122; 75% contribution from Saqqaq) (Figure 4, Table S4). These results reveal shared Siberian ancestry between ancient North Athabaskan and present-day Yeniseian populations.

## Discussion

In this study, we reconstruct detailed population dynamics of ancient Siberian populations and propose the Neolithic Yakutia populations as the origin of the Siberian ancestry. While ancient individuals with the APS ancestry are sporadically found across Siberia from Baikal to Arctic Siberia, it remains unclear how the APS populations related to each other and to later populations in the region (Sikora et al., 2019; Yu et al., 2020). By showing that two ancient individuals, irk030 and Dzhylinda-1, belong to the APS meta-population, we confirm a long-term presence of the APS population in southern Siberia between 14,000 and 6,000 years ago. Moreover, these two individuals represent two sub-lineages within the APS meta-population, forming the genetic substratum for the later Siberian populations in the West Baikal and Yakutia, respectively, thus suggesting that the divergence between the two regions dates back at least ca. 8,500 years ago. However, it is worth noting that this divergence does not mean a complete separation between the two regions, as shown by a gene flow from WestBaikal_EN to Yakutia_MN, indicating an expansion of Kitoi culture-related populations to the Yakutia region. Such a genetic connection aligns with archaeological studies (McKenzie, 2009; Kuzmin & Bellwood, 2015) describing the spread of net-impressed pottery style from west Baikal to Yakutia.

Contrary to previous suggestions of the Serovo-Glazkovo culture resulting from admixture between the preceding local Kitoi culture and an incoming ANE-rich population (de Barros Damgaard et al., 2018; Yu et al., 2020), our study presents findings that challenge this hypothesis. Specifically, our analysis of the earliest Serovo-Glazkovo individual, irk030, belongs to the APS ancestry and has no direct connection with the Kitoi culture-related populations. This suggests a clear genetic discontinuity in the west Baikal region consistent with a gap of archaeological records between the Kitoi and Serovo-Glazkovo cultures. Furthermore, our findings show that irk030 displays an expansion of APS ancestry into the Altai Mountains, thereby rejecting the notion of incoming ANE ancestry from Altai hunter-gatherers (Wang et al., 2023).

It is noteworthy that the Siberian ancestry found across Eurasia and North America can be traced to a single gene pool best represented by Yakutia_MN. The genetic difference between the Middle Neolithic Belkachi culture (Yakutia_MN) and Late Neolithic Ymyyakhtakh culture (Yakutia_LN) allows us to reason that the spread of the Siberian ancestry happened prior to Yakutia_LN. This fits with the time range given by the admixture date for the earliest presence of the Siberian ancestry in northeastern Europe (Bolshoy Oleni Ostrov; ca. 4,000 BP) and by the initial appearance of the Paleo-Eskimo culture (ca. 4,500 BP). This genetic evidence also aligns well with the dissemination of ceramic and lithic technologies, as documented by previous studies (Coutouly, 2016; Козлов et al., 2020). Interestingly, evidence of the second wave of Siberian ancestry expansion, associated with Ymyyakhtakh culture, is discernible in Nganasan individuals but not in another Samoyedic-speaking population, Selkup. This suggests that the divergence within Samoyedic-speaking populations may go back to the Neolithic period.

Despite our effort to construct a detailed historical model for the relationship between ancient and present-day Siberian populations, there are certain aspects that require further investigation. First, our understanding on the distribution and impact of the APS ancestry in Siberia is based only on a handful of ancient genomes, thus leaving it unknown how it became superseded by later migrant populations across Siberia. Second, Yakutia_MN and Russia_Bolshoy belong to different Y haplogroups, N and Q, respectively, which may be attributed to the limited availability of Middle Neolithic Yakutia genomes. We call for future paleogenomic studies to produce Middle Neolithic ancient genomes across Siberia, especially from Yakutia, to enhance our understanding on the details of the spread of the Siberian ancestry.

## Methods

### Genotype data preparation

We compiled previously published genome-wide genotype data of ancient individuals for the “1240K” panel, a set of 1,233,013 ancestry-informative single nucleotide polymorphisms (SNPs) (Mathieson et al., 2015; Fu et al., 2016). First, we took publicly available random pseudo-haploid pulldown genotype data for the 1240K panel from the Allen Ancient DNA Resource v37.2 and other individual studies (Table S1) (Rasmussen et al., 2010; Fu et al., 2014; Lazaridis et al., 2014; Raghavan et al., 2014; Rasmussen et al., 2014; Allentoft et al., 2015; Jones et al., 2015; Mathieson et al., 2015; Raghavan et al., 2015; Rasmussen et al., 2015; Fu et al., 2016; Jeong et al., 2016; Kılınç et al., 2016; Lazaridis et al., 2016; Lazaridis et al., 2017; Saag et al., 2017; Unterländer et al., 2017; Yang et al., 2017; Harney et al., 2018; Jeong et al., 2018; Lipson et al., 2018; Mathieson et al., 2018; Mittnik et al., 2018; Moreno-Mayar et al., 2018; Narasimhan et al., 2019; Jeong et al., 2020; Ning et al., 2020; Skourtanioti et al., 2020; Yang et al., 2020; Yu et al., 2020; Wang et al., 2021). Second, for ancient individuals whose genotype data are not available but aligned reads (BAMs) are, we obtained BAM files from the European Nucleotide Archive (https://www.ebi.ac.uk/ena), using the accession numbers provided in the original publications (Damgaard et al., 2018; de Barros Damgaard et al., 2018; Krzewińska et al., 2018; Lamnidis et al., 2018; Sikora et al., 2019; Kılınç et al., 2021; Wang et al., 2023). Table S1 provides a summary of the reference and raw data type (i.e., 1240K pull-down haploid genotype or BAM) for each ancient genome data.

Before genotyping with BAM file, we eliminated polymerase chain reaction duplicates using dedup v0.12.8 (Peltzer et al., 2016), and removed low quality reads with Phred-scaled mapping quality score lower than 30 using SAMtools v1.9 (Li et al., 2009). We then examined the pattern of post-mortem chemical damages using mapDamage program v2.2.1 (Jónsson et al., 2013), to ensure that it matched the expected pattern from the reported library preparation method. To minimize the impact of chemical damages in genotyping, we trimmed 3 and 10 bps at both ends of reads for non-UDG and partial-UDG treated double-strand libraries, respectively, using bamUtils v1.0.15 (Jun et al., 2015).

Using these BAM files, we produced random pseudo-haploid genotype data for the 1240K panel by randomly choosing one high-quality base (Phred-scaled base quality score 30 or higher) using the pileupCaller v1.5.2 with “randomHaploid” option (https://github.com/stschiff/sequenceTools; v1.5.2 last accessed at 19 April 2023). For double-stranded library data, we used ends-masked BAM files for transition SNPs and non-masked BAM files for transversions. For single-stranded library data, we used non-masked BAM files and applied the “singleStrandMode” option: this minimizes the impact of chemical damages by disregarding forward strand reads for C/T SNPs and reverse strand reads for G/A SNPs. We intersected the 1240K genotype data of ancient individuals with two sets of present-day worldwide individuals: 1) 1240K genotype data of individuals from the Simons Genome Diversity Project (Mallick et al., 2016), and 2) a broader set of individuals genotyped on the Affymetrix Axiome Genome-wide Human Origins 1 (”HumanOrigins”) (Patterson et al., 2012; Lazaridis et al., 2016; Flegontov et al., 2017; Jeong et al., 2019). For allele frequency-based analyses, our primary dataset of choice was the 1240K set. However, we utilized the HumanOrigins dataset for PCA and allele frequency-based analyses in cases where present-day Nganasan, Ket, and Selkup populations were included.

We extracted meta information about key ancient genome data including latitude, longitude, sex, radiocarbon date, haplogroup and relatedness from Allen Ancient DNA Resource (AADR) v52.2 (Mallick et al., 2023). For each first-degree pair or duplicate, we removed one of the individuals with lower sequencing coverage for further analysis.

### Principal components analysis (PCA)

We conducted principal components analysis (PCA) with present-day individuals genotyped on HumanOrigins using *smartpca* v18140 from EIGENSOFT v8.0.0 (Patterson et al., 2006). We used two population sets, the first including present-day Eurasian and American (2,270) and the second including Eurasian only (2,077). We projected ancient individuals not included in the PC calculation using the ‘lsqrproject: YES’ option. Samples used in PCA are listed in Table S1.

### *f*-statistics

We calculated the *f*-statistics by *qp3pop* and *f₄* functions from the R library ADMIXTOOLS2 v2.0.0. (https://github.com/uqrmaie1/admixtools, publication pending). We calculated outgroup-*fз* using the central African population Mbuti as an outgroup to measure shared genetic drift between target populations. Likewise, Mbuti was used as an outgroup to calculate *f₄* statistics in the form of *f₄*(Mbuti, X; target1, target2) for testing symmetricity between targets or searching additional admixture sources. Populations used in *f*-statistics are listed in Table S1, and the results of *f*-statistics are summarized in Table S5 and S6.

### qpWave and qpAdm analysis

We used *qpwave* and *qpadm* functions from the R library ADMIXTOOLS2 v2.0.0. for admixture modeling analysis. We used the following populations as a base outgroup set for both *qpWave* and *qpAdm* analysis: present-day Central African hunter-gatherers Mbuti (1240K: n=5; HumanOrigins: n=10), Taiwanese Aborigines Ami (1240K: n=2; HumanOrigins: n=10), Native Americans Mixe (1240K: n=5; HumanOrigins: n=10), indigenous Andamanese islander Onge (1240K: n=2; HumanOrigins: n=11), early Neolithic Iranians from the Ganj Dareh site Iran_N (n=8) (Lazaridis et al., 2016; Narasimhan et al., 2019), Epipaleolithic European Villabruna (n=1) (Fu et al., 2016), early Neolithic farmers from western Anatolia Anatolia_N (n=23) (Mathieson et al., 2015), and Neolithic southern Russia West_Siberia_N (n=3) (Narasimhan et al., 2019). In addition, when multiple admixture models were feasible, *qpAdm* rotating approach, which systematically shifts candidates from source to outgroup, was used to find the best proximal source.

### Graph-based analysis

In order to test component-wise admixture models, graph-based analysis was implemented using the *qpgraph* function from the R library ADMIXTOOLS2 v2.0.0. Before graph fitting, *f₂* statistics between all pairs of targets were calculated by the *extract_f2* function in ADMIXTOOLS2 with the ‘max_miss=0’ option, the same as the ‘allsnps: NO’ option from the previous version. The number of SNPs remaining by applying this option was 182,628. Mbuti population was also used as an outgroup in this analysis, and the following populations were used for distal representatives: MA1 for ANE; WHG for Mesolithic hunter-gatherers from Europe; USR1 for Native Americans; EastBaikal_N for ANA. Then, Middle Holocene populations were systematically added by following orders: irk030, Dzhylinda-1, WestBaikal_EN, Yakutia_MN, Saqqaq, WestBaikal_LNBA, and Yakutia_LN. The estimated branch length and admixture proportions were converted to dot file by in-house code, and we plotted admixture graph using Graphviz 6.0.1.

## Data Availability

Allen Ancient DNA Resource (AADR) v37.2 datasets were derived from sources in the public domain: https://reich.hms.harvard.edu/allen-ancient-dna-resource-aadr-downloadable-genotypes-present-day-and-ancient-dna-data, and v52.2 annotation information were derived from sources in the public domain: https://reich.hms.harvard.edu/allen-ancient-dna-resource-aadr-downloadable-genotypes-present-day-and-ancient-dna-data. The public domain and accession numbers of other ancient genome data are listed in Table S1. Haploid genotype data of ancient genome used in this study for the 1240K panel will be made publicly available upon the publication of the manuscript.

## Code availability

All analyses performed in this study are based on publicly available programs. Program names, versions, and nondefault options are described in the “Methods” section. All scripts used for the analyses presented in this study will be made publicly available via the Github repository.

## Acknowledgements

This work was supported by National Research Foundation of Korea (NRF) grant funded by the Korea government (MSIT) RS-2023-00212640 (C.J.),

## Author contribution

C.J. conceived and supervised the study. H.G. and J.L. curated and analyzed data. C.J. and H.G. wrote the manuscript with the input from J.L..

## Reference

Allentoft, M. E., Sikora, M., Sjögren, K.-G., Rasmussen, S., Rasmussen, M., Stenderup, J., Damgaard, P. B., Schroeder, H., Ahlström, T., & Vinner, L. (2015). Population genomics of bronze age Eurasia. Nature, 522(7555), 167–172.

Coutouly, Y. A. G. (2016). Migrations and interactions in prehistoric Beringia: the evolution of Yakutian lithic technology. Antiquity, 90(349), 9–31.

Damgaard, P. d. B., Marchi, N., Rasmussen, S., Peyrot, M., Renaud, G., Korneliussen, T., Moreno-Mayar, J. V., Pedersen, M. W., Goldberg, A., & Usmanova, E. (2018). 137 ancient human genomes from across the Eurasian steppes. Nature, 557(7705), 369–374.

de Barros Damgaard, P., Martiniano, R., Kamm, J., Moreno-Mayar, J. V., Kroonen, G., Peyrot, M., Barjamovic, G., Rasmussen, S., Zacho, C., & Baimukhanov, N. (2018). The first horse herders and the impact of early Bronze Age steppe expansions into Asia. Science, 360(6396), eaar7711.

Flegontov, P., Altınışık, N. E., Changmai, P., Rohland, N., Mallick, S., Adamski, N., Bolnick, D. A., Broomandkhoshbacht, N., Candilio, F., & Culleton, B. J. (2019). Palaeo-Eskimo genetic ancestry and the peopling of Chukotka and North America. Nature, 570(7760), 236–240.

Flegontov, P., Altinişik, N. E., Changmai, P., Rohland, N., Mallick, S., Bolnick, D. A., Candilio, F., Flegontova, O., Jeong, C., & Harper, T. K. (2017). Paleo-Eskimo genetic legacy across North America. bioRxiv, 203018.

Fu, Q., Li, H., Moorjani, P., Jay, F., Slepchenko, S. M., Bondarev, A. A., Johnson, P. L., Aximu-Petri, A., Prüfer, K., & De Filippo, C. (2014). Genome sequence of a 45,000-year-old modern human from western Siberia. Nature, 514(7523), 445–449.

Fu, Q., Posth, C., Hajdinjak, M., Petr, M., Mallick, S., Fernandes, D., Furtwängler, A., Haak, W., Meyer, M., & Mittnik, A. (2016). The genetic history of ice age Europe. Nature, 534(7606), 200–205.

Harney, É ., May, H., Shalem, D., Rohland, N., Mallick, S., Lazaridis, I., Sarig, R., Stewardson, K., Nordenfelt, S., & Patterson, N. (2018). Ancient DNA from Chalcolithic Israel reveals the role of population mixture in cultural transformation. Nature communications, 9(1), 3336.

Harney, E., Patterson, N., Reich, D., & Wakeley, J. (2021). Assessing the performance of qpAdm: a statistical tool for studying population admixture. Genetics, 217(4), iyaa045.

Jeong, C., Balanovsky, O., Lukianova, E., Kahbatkyzy, N., Flegontov, P., Zaporozhchenko, V., Immel, A., Wang, C.-C., Ixan, O., & Khussainova, E. (2019). The genetic history of admixture across inner Eurasia. Nature ecology & evolution, 3(6), 966–976.

Jeong, C., Ozga, A. T., Witonsky, D. B., Malmström, H., Edlund, H., Hofman, C. A., Hagan, R. W., Jakobsson, M., Lewis, C. M., & Aldenderfer, M. S. (2016). Long-term genetic stability and a high-altitude East Asian origin for the peoples of the high valleys of the Himalayan arc. Proceedings of the National Academy of Sciences, 113(27), 7485–7490.

Jeong, C., Wang, K., Wilkin, S., Taylor, W. T. T., Miller, B. K., Bemmann, J. H., Stahl, R., Chiovelli, C., Knolle, F., & Ulziibayar, S. (2020). A dynamic 6,000-year genetic history of Eurasia’s Eastern Steppe. Cell, 183(4), 890–904. e829.

Jeong, C., Wilkin, S., Amgalantugs, T., Bouwman, A. S., Taylor, W. T. T., Hagan, R. W., Bromage, S., Tsolmon, S., Trachsel, C., & Grossmann, J. (2018). Bronze Age population dynamics and the rise of dairy pastoralism on the eastern Eurasian steppe. Proceedings of the National Academy of Sciences, 115(48), E11248–E11255.

Jones, E. R., Gonzalez-Fortes, G., Connell, S., Siska, V., Eriksson, A., Martiniano, R., McLaughlin, R. L., Gallego Llorente, M., Cassidy, L. M., & Gamba, C. (2015). Upper Palaeolithic genomes reveal deep roots of modern Eurasians. Nature communications, 6(1), 8912.

Jónsson, H., Ginolhac, A., Schubert, M., Johnson, P. L., & Orlando, L. (2013). mapDamage2. 0: fast approximate Bayesian estimates of ancient DNA damage parameters. bioinformatics, 29(13), 1682–1684.

Jun, G., Wing, M. K., Abecasis, G. R., & Kang, H. M. (2015). An efficient and scalable analysis framework for variant extraction and refinement from population-scale DNA sequence data. Genome research, 25(6), 918–925.

Karafet, T. M., Osipova, L. P., Savina, O. V., Hallmark, B., & Hammer, M. F. (2018). Siberian genetic diversity reveals complex origins of the Samoyedic-speaking populations. American Journal of Human Biology, 30(6), e23194.

Kılınç, G. M., Kashuba, N., Koptekin, D., Bergfeldt, N., Dönertaş, H. M., Rodríguez-Varela, R., Shergin, D., Ivanov, G., Kichigin, D., & Pestereva, K. (2021). Human population dynamics and Yersinia pestis in ancient northeast Asia. Science advances, 7(2), eabc4587.

Kılınç, G. M., Omrak, A., Özer, F., Günther, T., Büyükkarakaya, A. M., Bıçakçı, E., Baird, D., Dönertaş, H. M., Ghalichi, A., & Yaka, R. (2016). The demographic development of the first farmers in Anatolia. Current Biology, 26(19), 2659–2666.

Krzewińska, M., Kılınç, G. M., Juras, A., Koptekin, D., Chyleński, M., Nikitin, A. G., Shcherbakov, N., Shuteleva, I., Leonova, T., & Kraeva, L. (2018). Ancient genomes suggest the eastern Pontic-Caspian steppe as the source of western Iron Age nomads. Science advances, 4(10), eaat4457.

Kuzmin, Y. V., & Bellwood, P. (2015). Northern and North-eastern Asia: archaeology. The global prehistory of human migration, 191–196.

Lamnidis, T. C., Majander, K., Jeong, C., Salmela, E., Wessman, A., Moiseyev, V., Khartanovich, V., Balanovsky, O., Ongyerth, M., & Weihmann, A. (2018). Ancient Fennoscandian genomes reveal origin and spread of Siberian ancestry in Europe. Nature communications, 9(1), 1–12.

Lazaridis, I., Mittnik, A., Patterson, N., Mallick, S., Rohland, N., Pfrengle, S., Furtwängler, A., Peltzer, A., Posth, C., & Vasilakis, A. (2017). Genetic origins of the Minoans and Mycenaeans. Nature, 548(7666), 214–218.

Lazaridis, I., Nadel, D., Rollefson, G., Merrett, D. C., Rohland, N., Mallick, S., Fernandes, D., Novak, M., Gamarra, B., & Sirak, K. (2016). Genomic insights into the origin of farming in the ancient Near East. Nature, 536(7617), 419–424.

Lazaridis, I., Patterson, N., Mittnik, A., Renaud, G., Mallick, S., Kirsanow, K., Sudmant, P. H., Schraiber, J. G., Castellano, S., & Lipson, M. (2014). Ancient human genomes suggest three ancestral populations for present-day Europeans. Nature, 513(7518), 409–413.

Li, H., Handsaker, B., Wysoker, A., Fennell, T., Ruan, J., Homer, N., Marth, G., Abecasis, G., Durbin, R., & Subgroup, G. P. D. P. (2009). The sequence alignment/map format and SAMtools. bioinformatics, 25(16), 2078–2079.

Lipson, M., Cheronet, O., Mallick, S., Rohland, N., Oxenham, M., Pietrusewsky, M., Pryce, T. O., Willis, A., Matsumura, H., & Buckley, H. (2018). Ancient genomes document multiple waves of migration in Southeast Asian prehistory. Science, 361(6397), 92–95.

Mallick, S., Li, H., Lipson, M., Mathieson, I., Gymrek, M., Racimo, F., Zhao, M., Chennagiri, N., Nordenfelt, S., & Tandon, A. (2016). The Simons genome diversity project: 300 genomes from 142 diverse populations. Nature, 538(7624), 201–206.

Mallick, S., Micco, A., Mah, M., Ringbauer, H., Lazaridis, I., Olalde, I., Patterson, N. J., & Reich, D. E. (2023). The Allen Ancient DNA Resource (AADR): A curated compendium of ancient human genomes. bioRxiv, 2023.2004. 2006.535797.

Mao, X., Zhang, H., Qiao, S., Liu, Y., Chang, F., Xie, P., Zhang, M., Wang, T., Li, M., & Cao, P. (2021). The deep population history of northern East Asia from the Late Pleistocene to the Holocene. Cell, 184(12), 3256–3266. e3213.

Mathieson, I., Alpaslan-Roodenberg, S., Posth, C., Szécsényi-Nagy, A., Rohland, N., Mallick, S., Olalde, I., Broomandkhoshbacht, N., Candilio, F., & Cheronet, O. (2018). The genomic history of southeastern Europe. Nature, 555(7695), 197–203.

Mathieson, I., Lazaridis, I., Rohland, N., Mallick, S., Patterson, N., Roodenberg, S. A., Harney, E., Stewardson, K., Fernandes, D., & Novak, M. (2015). Genome-wide patterns of selection in 230 ancient Eurasians. Nature, 528(7583), 499–503.

McKenzie, H. G. (2009). Review of early hunter-gatherer pottery in Eastern Siberia. In M. Z. Peter Jordan (Ed.), Ceramics before Farming: The Dispersal of Pottery among Prehistoric Eurasian Hunter-Gatherers (pp. 167–208). Walnut Creek: Left Coast Press.

Mittnik, A., Wang, C.-C., Pfrengle, S., Daubaras, M., Zariņa, G., Hallgren, F., Allmäe, R., Khartanovich, V., Moiseyev, V., & Tõrv, M. (2018). The genetic prehistory of the Baltic Sea region. Nature communications, 9(1), 442.

Moreno-Mayar, J. V., Potter, B. A., Vinner, L., Steinrücken, M., Rasmussen, S., Terhorst, J., Kamm, J. A., Albrechtsen, A., Malaspinas, A.-S., & Sikora, M. (2018). Terminal Pleistocene Alaskan genome reveals first founding population of Native Americans. Nature, 553(7687), 203–207.

Movsesian, A. A., Bakholdina, V. Y., & Pezhemsky, D. V. (2014). Biological diversity and population history of M iddle H olocene hunter-gatherers from the C is-B aikal region of S iberia. American Journal of Physical Anthropology, 155(4), 559–570.

Narasimhan, V. M., Patterson, N., Moorjani, P., Rohland, N., Bernardos, R., Mallick, S., Lazaridis, I., Nakatsuka, N., Olalde, I., & Lipson, M. (2019). The formation of human populations in South and Central Asia. Science, 365(6457), eaat7487.

Ning, C., Li, T., Wang, K., Zhang, F., Li, T., Wu, X., Gao, S., Zhang, Q., Zhang, H., & Hudson, M. J. (2020). Ancient genomes from northern China suggest links between subsistence changes and human migration. Nature communications, 11(1), 1–9.

Patterson, N., Moorjani, P., Luo, Y., Mallick, S., Rohland, N., Zhan, Y., Genschoreck, T., Webster, T., & Reich, D. (2012). Ancient admixture in human history. Genetics, 192(3), 1065–1093.

Patterson, N., Price, A. L., & Reich, D. (2006). Population structure and eigenanalysis. PLoS genetics, 2(12), e190.

Peltola, S., Majander, K., Makarov, N., Dobrovolskaya, M., Nordqvist, K., Salmela, E., & Onkamo, P. (2023). Genetic admixture and language shift in the medieval Volga-Oka interfluve. Current Biology, 33(1), 174–182. e110.

Peltzer, A., Jäger, G., Herbig, A., Seitz, A., Kniep, C., Krause, J., & Nieselt, K. (2016). EAGER: efficient ancient genome reconstruction. Genome Biology, 17(1), 1–14.

Powers, W. R., & Jordan, R. H. (1990). Human biogeography and climate change in Siberia and Arctic North America in the fourth and fifth millennia BP. *Philosophical Transactions of the Royal Society of London. Series A*, Mathematical and Physical Sciences, 330(1615), 665–670.

Raghavan, M., Skoglund, P., Graf, K. E., Metspalu, M., Albrechtsen, A., Moltke, I., Rasmussen, S., Stafford Jr, T. W., Orlando, L., & Metspalu, E. (2014). Upper Palaeolithic Siberian genome reveals dual ancestry of Native Americans. Nature, 505(7481), 87–91.

Raghavan, M., Steinrücken, M., Harris, K., Schiffels, S., Rasmussen, S., DeGiorgio, M., Albrechtsen, A., Valdiosera, C., Ávila-Arcos, M. C., & Malaspinas, A.-S. (2015). Genomic evidence for the Pleistocene and recent population history of Native Americans. Science, 349(6250), aab3884.

Rasmussen, M., Anzick, S. L., Waters, M. R., Skoglund, P., DeGiorgio, M., Stafford, T. W., Rasmussen, S., Moltke, I., Albrechtsen, A., & Doyle, S. M. (2014). The genome of a Late Pleistocene human from a Clovis burial site in western Montana. Nature, 506(7487), 225–229.

Rasmussen, M., Li, Y., Lindgreen, S., Pedersen, J. S., Albrechtsen, A., Moltke, I., Metspalu, M., Metspalu, E., Kivisild, T., & Gupta, R. (2010). Ancient human genome sequence of an extinct Palaeo-Eskimo. Nature, 463(7282), 757–762.

Rasmussen, M., Sikora, M., Albrechtsen, A., Korneliussen, T. S., Moreno-Mayar, J. V., Poznik, G. D., Zollikofer, C. P., Ponce de León, M. S., Allentoft, M. E., & Moltke, I. (2015). The ancestry and affiliations of Kennewick Man. Nature, 523(7561), 455–458.

Reich, D., Patterson, N., Campbell, D., Tandon, A., Mazieres, S., Ray, N., Parra, M. V., Rojas, W., Duque, C., & Mesa, N. (2012). Reconstructing native American population history. Nature, 488(7411), 370–374.

Saag, L., Laneman, M., Varul, L., Malve, M., Valk, H., Razzak, M. A., Shirobokov, I. G., Khartanovich, V. I., Mikhaylova, E. R., & Kushniarevich, A. (2019). The arrival of Siberian ancestry connecting the Eastern Baltic to Uralic speakers further East. Current Biology, 29(10), 1701–1711. e1716.

Saag, L., Varul, L., Scheib, C. L., Stenderup, J., Allentoft, M. E., Saag, L., Pagani, L., Reidla, M., Tambets, K., & Metspalu, E. (2017). Extensive farming in Estonia started through a sex-biased migration from the Steppe. Current Biology, 27(14), 2185–2193. e2186.

Sandweiss, D. H., Maasch, K. A., & Anderson, D. G. (1999). Transitions in the mid-Holocene. Science, 283(5401), 499–500.

Sikora, M., Pitulko, V. V., Sousa, V. C., Allentoft, M. E., Vinner, L., Rasmussen, S., Margaryan, A., de Barros Damgaard, P., de la Fuente, C., & Renaud, G. (2019). The population history of northeastern Siberia since the Pleistocene. Nature, 570(7760), 182–188.

Siska, V., Jones, E. R., Jeon, S., Bhak, Y., Kim, H.-M., Cho, Y. S., Kim, H., Lee, K., Veselovskaya, E., & Balueva, T. (2017). Genome-wide data from two early Neolithic East Asian individuals dating to 7700 years ago. Science advances, 3(2), e1601877.

Skourtanioti, E., Erdal, Y. S., Frangipane, M., Restelli, F. B., Yener, K. A., Pinnock, F., Matthiae, P., Özbal, R., Schoop, U.-D., & Guliyev, F. (2020). Genomic history of neolithic to bronze age Anatolia, northern Levant, and southern Caucasus. Cell, 181(5), 1158–1175. e1128.

Tambets, K., Yunusbayev, B., Hudjashov, G., Ilumäe, A.-M., Rootsi, S., Honkola, T., Vesakoski, O., Atkinson, Q., Skoglund, P., & Kushniarevich, A. (2018). Genes reveal traces of common recent demographic history for most of the Uralic-speaking populations. Genome Biology, 19(1), 1–20.

Unterländer, M., Palstra, F., Lazaridis, I., Pilipenko, A., Hofmanová, Z., Groß, M., Sell, C., Blöcher, J., Kirsanow, K., & Rohland, N. (2017). Ancestry and demography and descendants of Iron Age nomads of the Eurasian Steppe. Nature communications, 8(1), 14615.

Wang, C.-C., Yeh, H.-Y., Popov, A. N., Zhang, H.-Q., Matsumura, H., Sirak, K., Cheronet, O., Kovalev, A., Rohland, N., & Kim, A. M. (2021). Genomic insights into the formation of human populations in East Asia. Nature, 591(7850), 413–419.

Wang, K., Yu, H., Radzevičiūtė, R., Kiryushin, Y. F., Tishkin, A. A., Frolov, Y. V., Stepanova, N. F., Kiryushin, K. Y., Kungurov, A. L., & Shnaider, S. V. (2023). Middle Holocene Siberian genomes reveal highly connected gene pools throughout North Asia. Current Biology.

Weber, A. W., Link, D. W., & Katzenberg, M. A. (2002). Hunter-gatherer culture change and continuity in the Middle Holocene of the Cis-Baikal, Siberia. Journal of Anthropological Archaeology, 21(2), 230–299.

Yang, M. A., Fan, X., Sun, B., Chen, C., Lang, J., Ko, Y.-C., Tsang, C.-h., Chiu, H., Wang, T., & Bao, Q. (2020). Ancient DNA indicates human population shifts and admixture in northern and southern China. Science, 369(6501), 282–288.

Yang, M. A., Gao, X., Theunert, C., Tong, H., Aximu-Petri, A., Nickel, B., Slatkin, M., Meyer, M., Pääbo, S., & Kelso, J. (2017). 40,000-year-old individual from Asia provides insight into early population structure in Eurasia. Current Biology, 27(20), 3202–3208. e3209.

Yu, H., Spyrou, M. A., Karapetian, M., Shnaider, S., Radzevičiūtė, R., Nägele, K., Neumann, G. U., Penske, S., Zech, J., & Lucas, M. (2020). Paleolithic to Bronze Age Siberians reveal connections with first Americans and across Eurasia. Cell, 181(6), 1232–1245. e1220.

Козлов, А. И., Вершубская, Г. Г., & Боринская, С. А. (2020). Дивергенция генетических комплексов у антропологически родственных популяций при разных типах хозяйствования. Вестник Московского университета. Серия 23. *Антропология*(4), 99–110.

