## Supplementary figures for "Reconstructing the genetic relationship between ancient and present-day Siberian populations"

**This file includes:**

Supplementary Figures S1 to S5

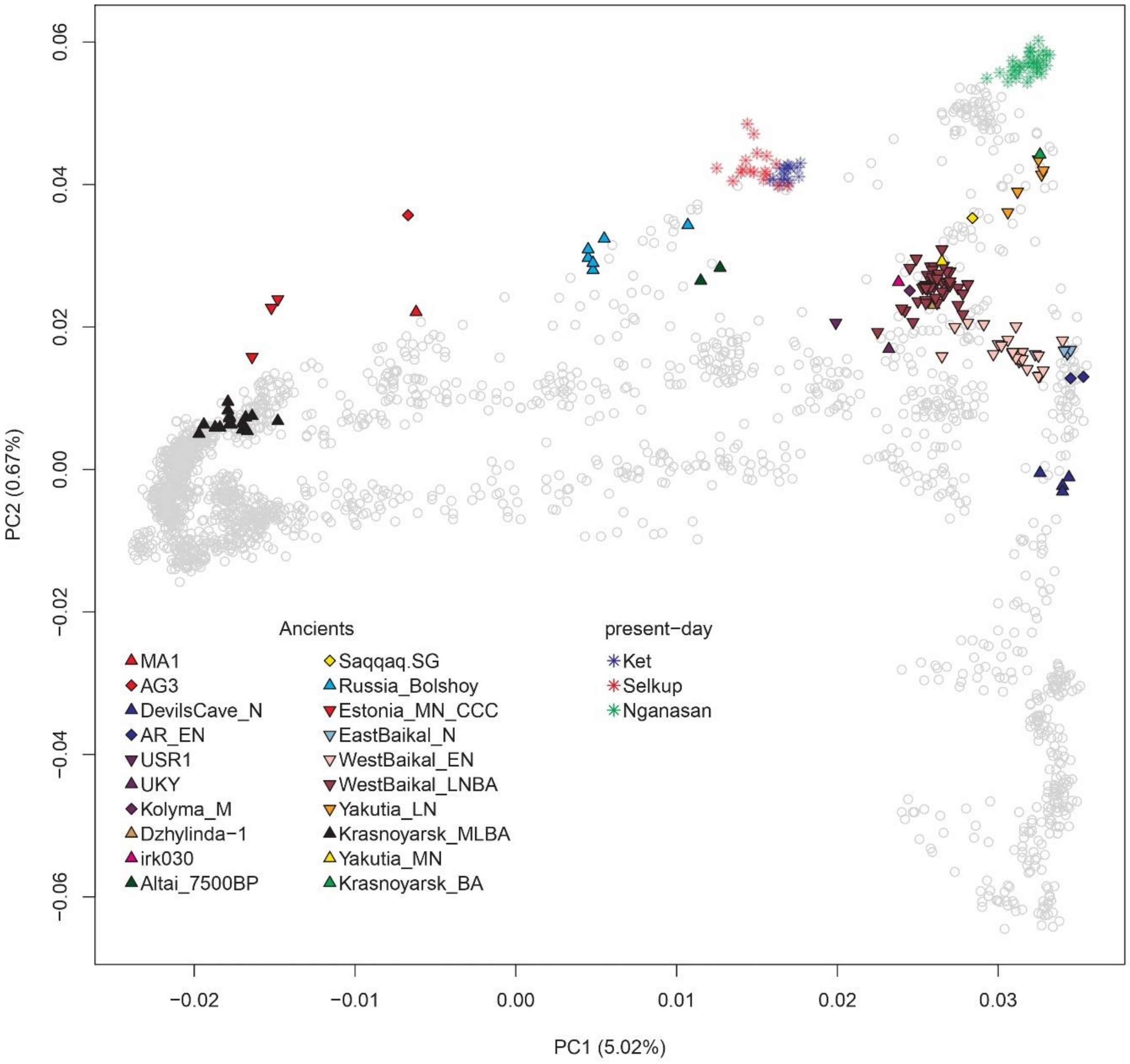


**Figure S1. Principal component analysis performed with present-day Eurasian individuals.** The principal component analysis is performed with present-day Eurasian individuals, and each present-day sample is placed on principal component 1 and 2 coordinates by grey circles. Key ancient and present-day genomes are projected on pre-calculated principal components and labeled.

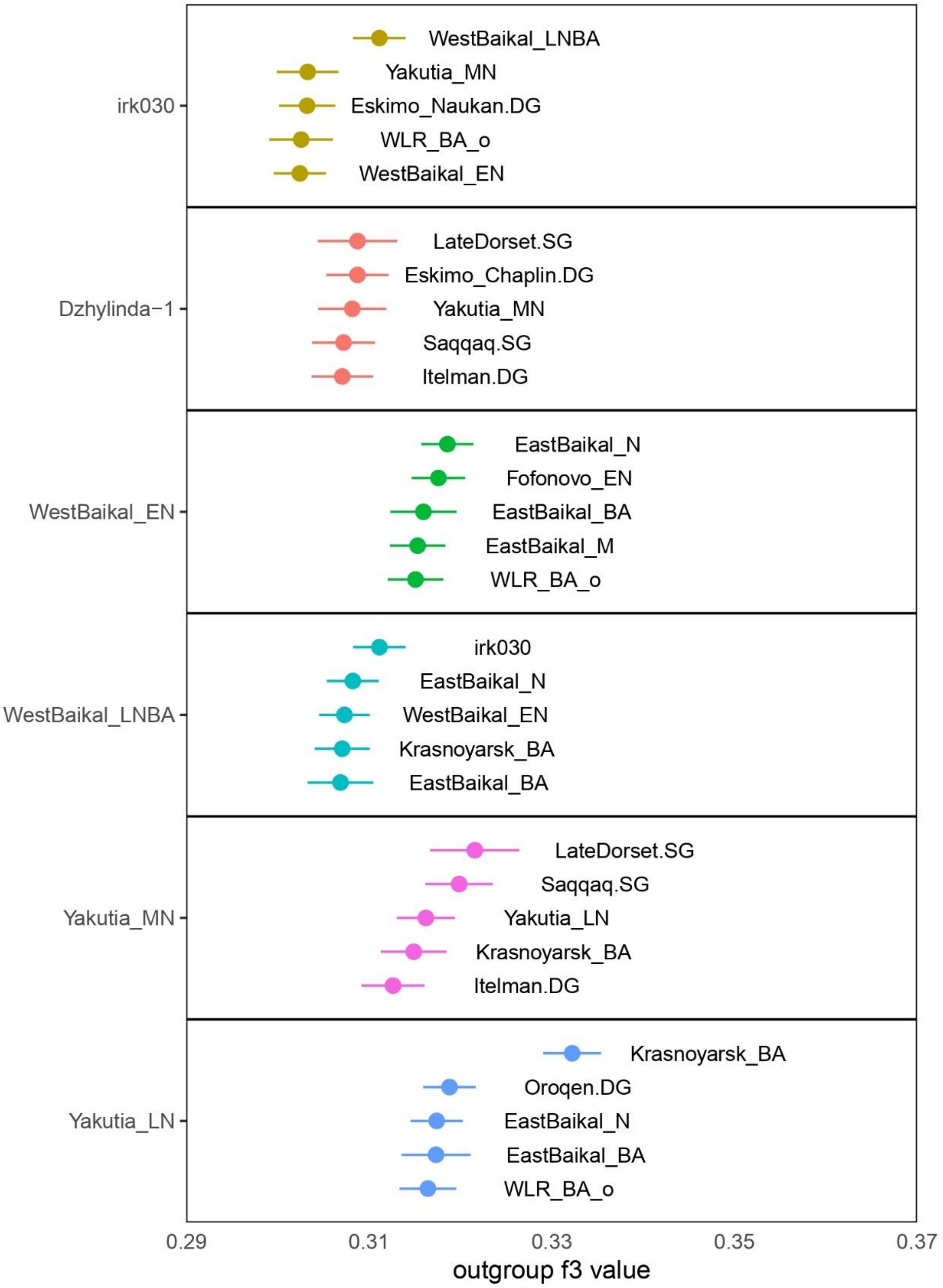


**Figure S2. Outgroup-$f_3$ values for the Middle Holocene Siberian populations.** Outgroup-$f_3$ statistics are calculated in the form of $f_3$(Mbuti; X, Target), and the highest 5 populations for each target are shown. Horizontal bars represent the point estimate ±1 SE, and SEs are calculated by 5cM block jackknifing. All outgroup-$f_3$ values are reported in Table S6.

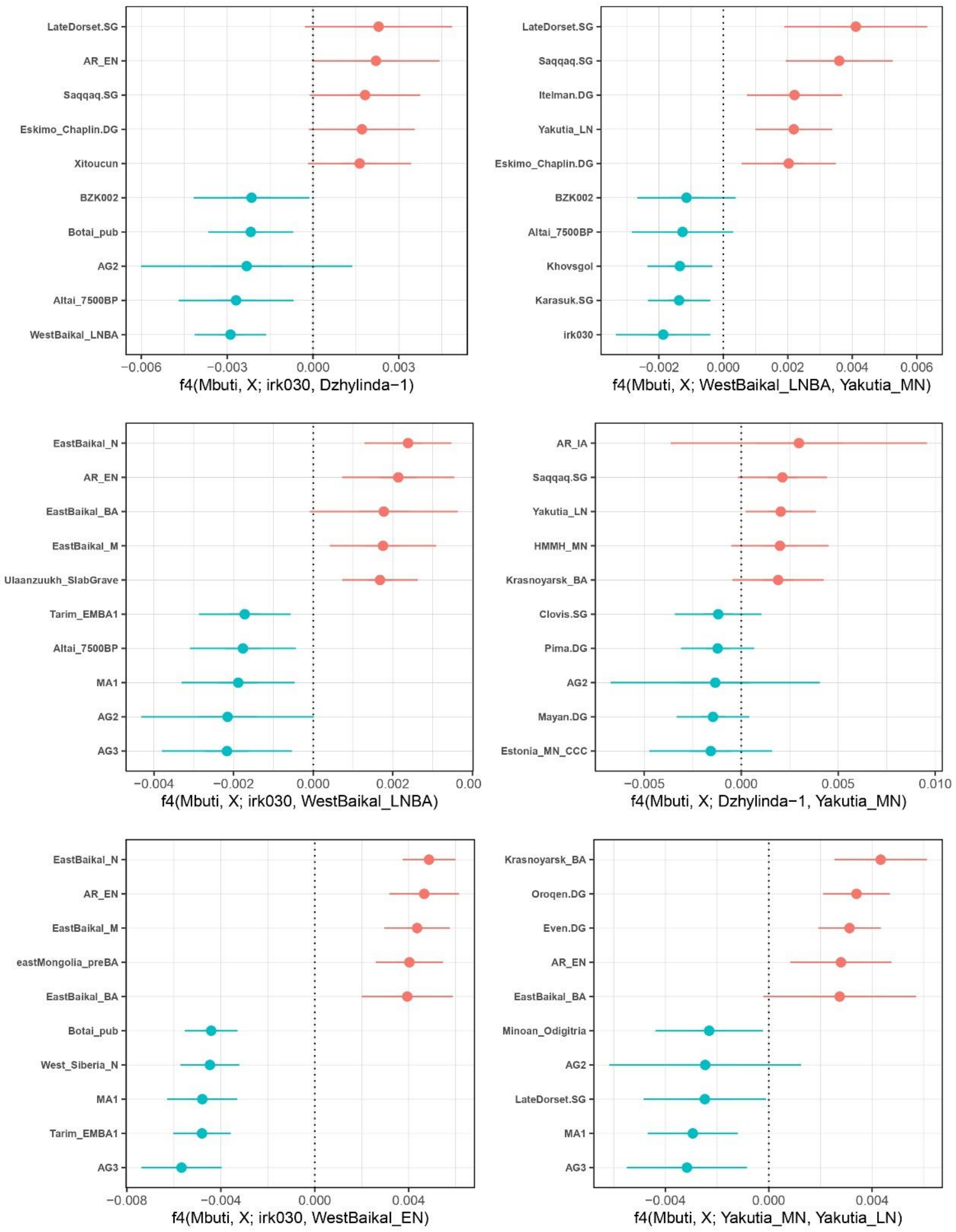


**Figure S3. Genetic symmetricity test between Middle Holocene Siberians.** $F_4$ statistics in the form $f_4$(Mbuti, X; target1, target2) are calculated. The 5 most positive and 5 most negative $f_4$ statistics are shown in red and blue, respectively. Horizontal bars represent the point estimate ±3 SE, respectively. SEs are calculated by 5cM block jackknifing. All $f_4$ values are reported in Table S5.

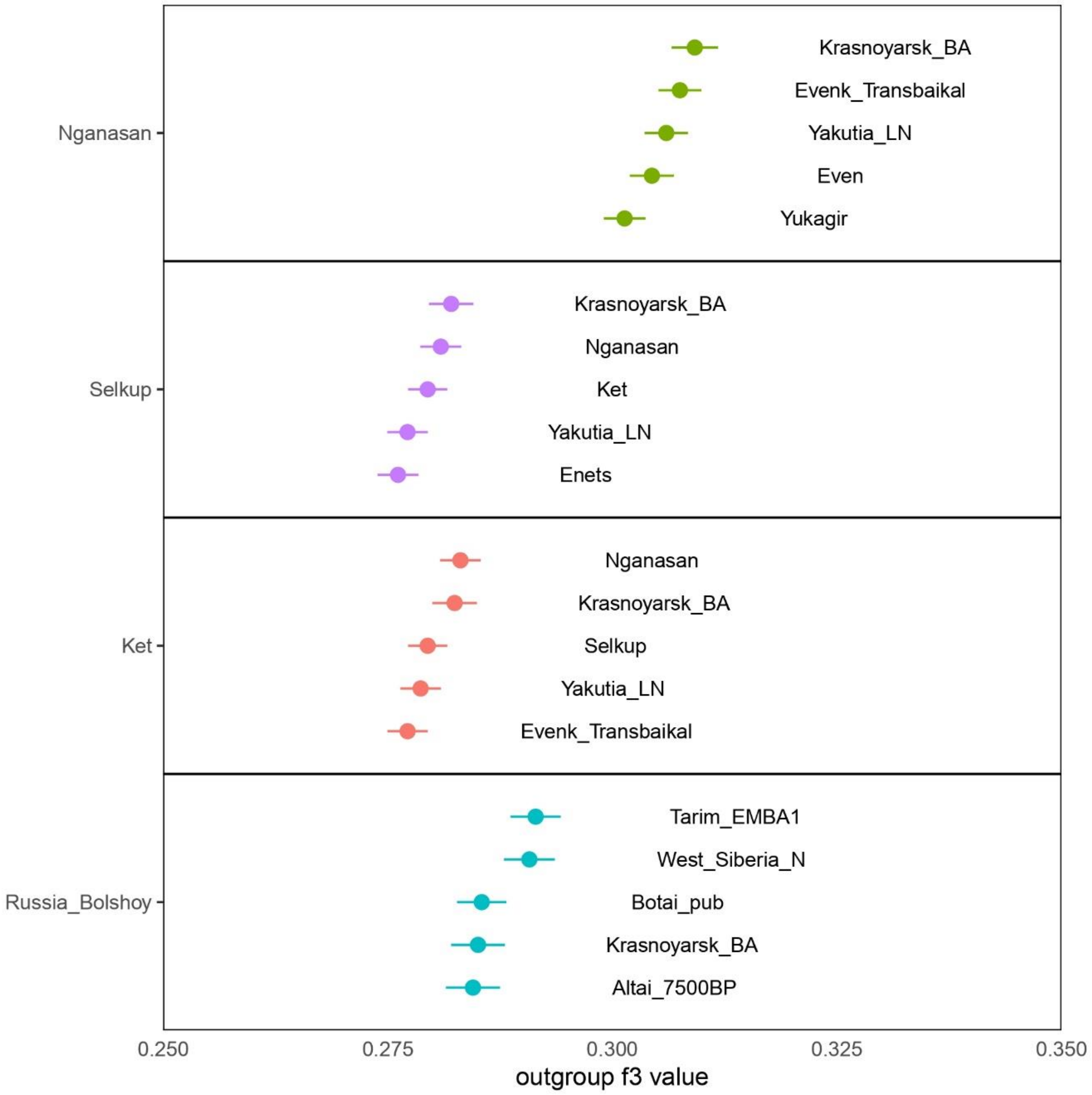


**Figure S4. Outgroup-$f_3$ values for the Siberian ancestry-related populations.** Outgroup-$f_3$ statistics are calculated in the form of $f_3$(Mbuti; X, Target), and the highest 5 populations for each target are shown. Horizontal bars represent the point estimate ±1 standard error, and standard errors are calculated by 5cM block jackknifing. All outgroup-$f_3$ values are reported in Table S6.

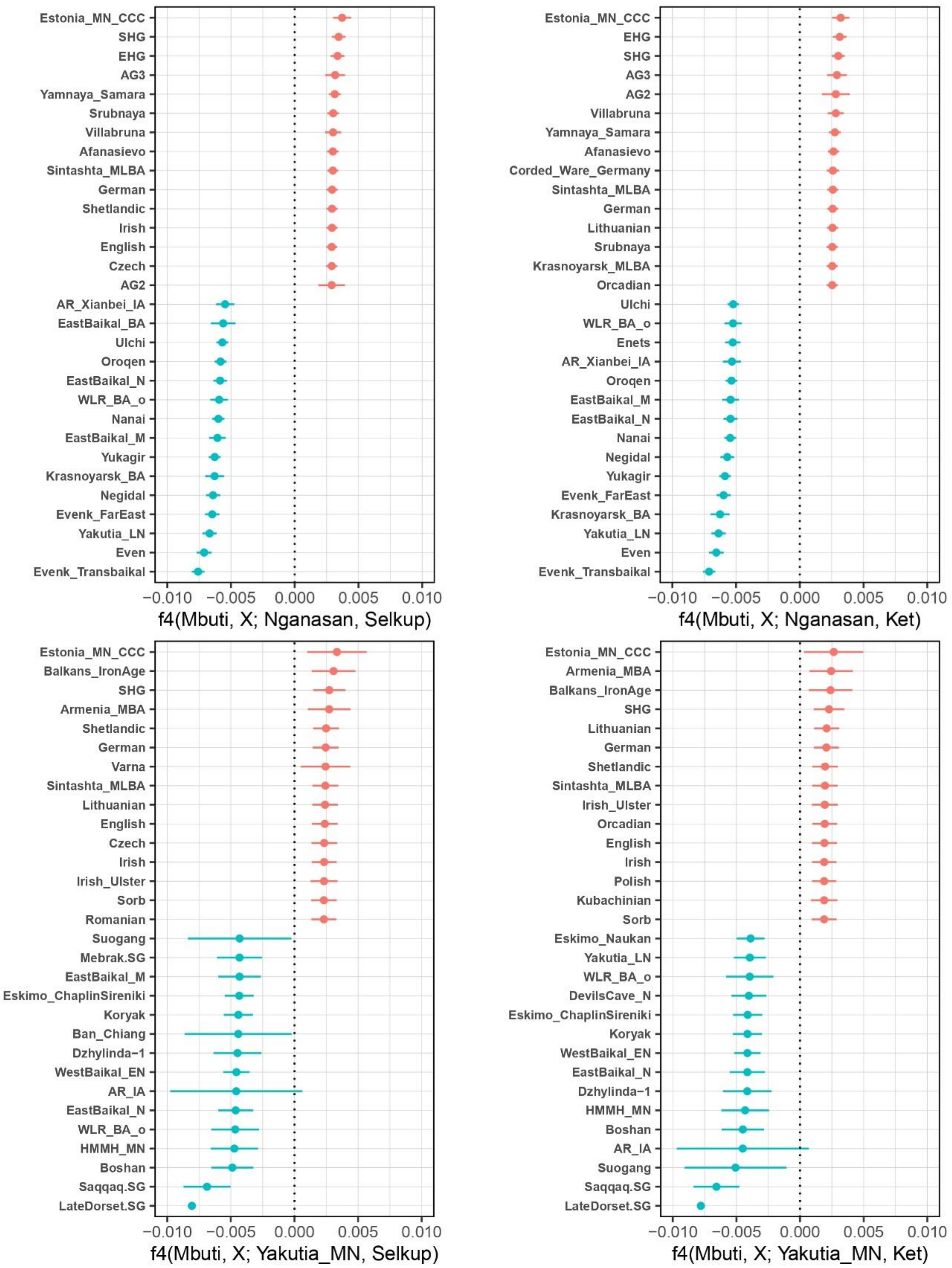


**Figure S5. Genetic symmetricity test for present-day Selkup and Ket populations.** $F_4$ statistics in the form $f_4$(Mbuti, X; target1, target2) are calculated. The 5 most positive and 5 most negative $f_4$ statistics are shown in red and blue, respectively. Horizontal bars represent the point estimate ±3 SE, respectively. SEs are calculated by 5cM block jackknifing. All $f_4$ values are reported in Table S5.
